# GNN2Pfam: Integrating protein sequence and structure with graph neural networks for Pfam domain annotation

**DOI:** 10.1101/2025.09.18.677074

**Authors:** Emilio Fenoy, Leandro A. Bugnon, Rosario Vitale, Sofia A. Duarte, Diego H. Milone, Georgina Stegmayer

## Abstract

The challenge of establishing the relationship between protein sequences and their function cannot yet be considered completely solved. State-of-the-art annotation of Pfam domains is based on hidden Markov models (HMMs) built from hand-crafted sequence alignments. However, while this approach has been highly successful during the last decades since its proposal, there is yet a very large number of proteins that remain unannotated because there is no possible alignment to already known and functionally characterized sequences, or HMM fails to discriminate between similar domains. Adding structural information using deep and graph neural networks (GNNs) presents an opportunity to build upon existing models in those more challenging cases. GNNs excel at capturing complex relationships in data and can learn a model that shares information across all existing families, thus being able to generalize Pfam domain predictions to novel sequences. In this work we propose GNN2Pfam, an end-to-end GNN-based method for Pfam family domain annotation. Our strategy allows one single model to be trained for all species and families. This novel proposal uses the protein 3D structure together with a sequence representation obtained from a large pre-trained model. The GNN2Pfam model is based on a graph derived from amino acid interactions in the 3D structure, learning both sequential and structural features from this representation. Experiments show that the proposed GNN-based model can clearly outperform the HMM state-of-the-art predictive performance in Pfam domains annotations. These results suggest that GNN models can be the key component of future protein annotation tools. Data and source code are available at https://github.com/efenoy/GNN2Pfam.

## 1. Introduction

Proteins are building blocks of life, playing many crucial roles within organisms, such as catalyzing chemical reactions, coordinating signal pathways and providing structural support to cells (28). In order to elucidate the mechanism of life, it is important to identify protein functions, which are closely related to their domains. Domains are distinct functional and/or structural units in a protein. Usually they are responsible for a particular function or interaction, contributing to the overall role of a protein. Often each individual domain has a specific function, such as for example binding a particular molecule or catalysing a given reaction. Proteins are assigned to families according to the domain(s) they contain. A protein family is a group of proteins that share a common evolutionary origin, reflected by their related functions and similarities in sequence or structure.

The Pfam database is a comprehensive collection of protein domains and families used for protein structure and function analysis (18). Automated Pfam family prediction of proteins is a large-scale multi-label classification problem. Nowadays this task is performed through multiple sequence alignments and profile Hidden Markov Models (HMMs). Each Pfam family has a seed alignment containing a representative set of sequences, from which a profile HMM is generated. This HMM is then used to search against a database containing sequences from the UniProtKB reference proteomes using the HMMER software (11). Sequence regions that meet a family-specific curated threshold are aligned to the profile HMM to create a full alignment and the corresponding Pfam family annotation. This approach has some limitations: alignment based methods are not accurate enough; a single HMM model must be trained for each family, separately and independently; and HMMs are not fast enough to handle large numbers of protein sequences from numerous genomes. Thus, it is a current challenge to develop a powerful automatic annotation method for the proteins deposited in Pfam capable of overcoming these limitations.

In the last decade, deep learning (DL) has led to unprecedented improvements in a broad spectrum of problems, ranging from learning protein sequence embeddings to predicting protein structure (27; 13) and function (15; 16). In recent years DL methods for modeling Pfam protein families appeared (21; 2), using only sequence information and with the capability of learning from a complete dataset, thus being able to discover inner patterns across several families at the same time. Models can be fit from scratch (15; 2; 4) or fine-tuned from a model pretrained on unlabeled protein sequences (22; 17; 31) since the amino acid sequence largely specifies a protein structure and function (1).

Recent work has shown that DL models for protein Pfam functional predictions together with transfer learning and representations obtained from a protein large language model (pLM) can effectively outperform traditional techniques (26). In fact several pre-trained pLMs have appeared in the last five years (30; 7; 23) that take advantage of the vast quantity of unannotated protein sequence data available. A pre-trained pLM, given a raw protein sequence, can calculate a low-dimensional feature vector (embedding) that encodes its representation. Next, a predictive model can solve a downstream task by learning from the embeddings that are associated with specific target labels. A review (10) where several protein sequence representation learning methods were experimentally benchmarked, indicated Evolutionary Scale Modeling (ESM) (20) as the best method for the tasks evaluated, which included Pfam family prediction.

Most sequence-based protein function prediction methods use multiple 1D convolutional neural network layers (CNNs) that search for spatial patterns within a given sequence and convert them into complex features using multiple convolutional layers. In contrast, graph neural networks (GNNs) have gained in popularity recently (3; 12) for structural protein function prediction since those can overcome these limitations by generalizing convolutional operations on more efficient graph-like representations (12; 24) and because those can learn transformed representations of interacting pairs of elements within a sequence via graph relationships. Precisely in the case of proteins and their domains, where 3D structure is of utmost importance for defining function, a GNN model can allow integrating not only sequential but also, and more important, structural information. GNNs are powerful deep-learning-based methods for learning rich context-informed representations of nodes in graphs by propagating and aggregating different types of input information (such as semantic and structural data representations) from a node and its local neighborhood (6). The transformed representations (embeddings) can be used then for several downstream tasks, such as Pfam domain classification. These methods have demonstrated great success in many biology and healthcare-related tasks (32; 29; 19).

In this work we propose GNN2Pfam, an end-to-end GNN-based method for Pfam family domain annotation. This novel proposal uses the protein 3D structure together with a sequence representation obtained from a large pre-trained model. The model is based on a graph derived from amino acid interactions in the 3D structure, learning both sequential and structural features from this representation. A last layer based on Conditional Random Fields (CRF) provides the output probabilities for each Pfam family along the sequence of amino acids. Our strategy allows one single model to be trained for all species and families. Extensive experiments with several datasets show that the proposed GNN-based approach clearly outperforms the HMM state-of-the-art method.

## GNN2Pfam: novel GNN for Pfam family annotation

We introduce a model that leverages graph-based representations by incorporating several GNN layers, which extract structural and relational features from the protein sequence and its structure. Figure 1 shows the pipeline of GNN2Pfam, the GNN model proposed for Pfam family annotation. For each input protein, its corresponding 3D structure is retrieved from the AlphaFold2 database (Fig. 1.I). Then, per-aminoacid embeddings are obtained from a pLM, and its corresponding 3D structure is used to build a graph derived from amino acid interactions in the structure (Fig. 1.II). Each node in the graph is the pLM embedding of an amino acid, and each edge has a 4×4 matrix of backbone atom distances. After that, the graph obtained feeds two GNN layers and a CRF layer that can take context into account and predict a family label, considering neighbouring samples as well (Fig. 1.III). Finally, at the output of the pipeline (Fig. 1.IV) the class scores are predicted for each amino acid in the sequence, obtaining a curve of scores for the final Pfam family prediction. The plot on the top of Fig. 1.IV shows the curve of scores for the winner predicted Pfam family (green line) and the no domain curve (red line); the corresponding 3D structure of the predicted domain is highlighted below.

**Figure 1:**
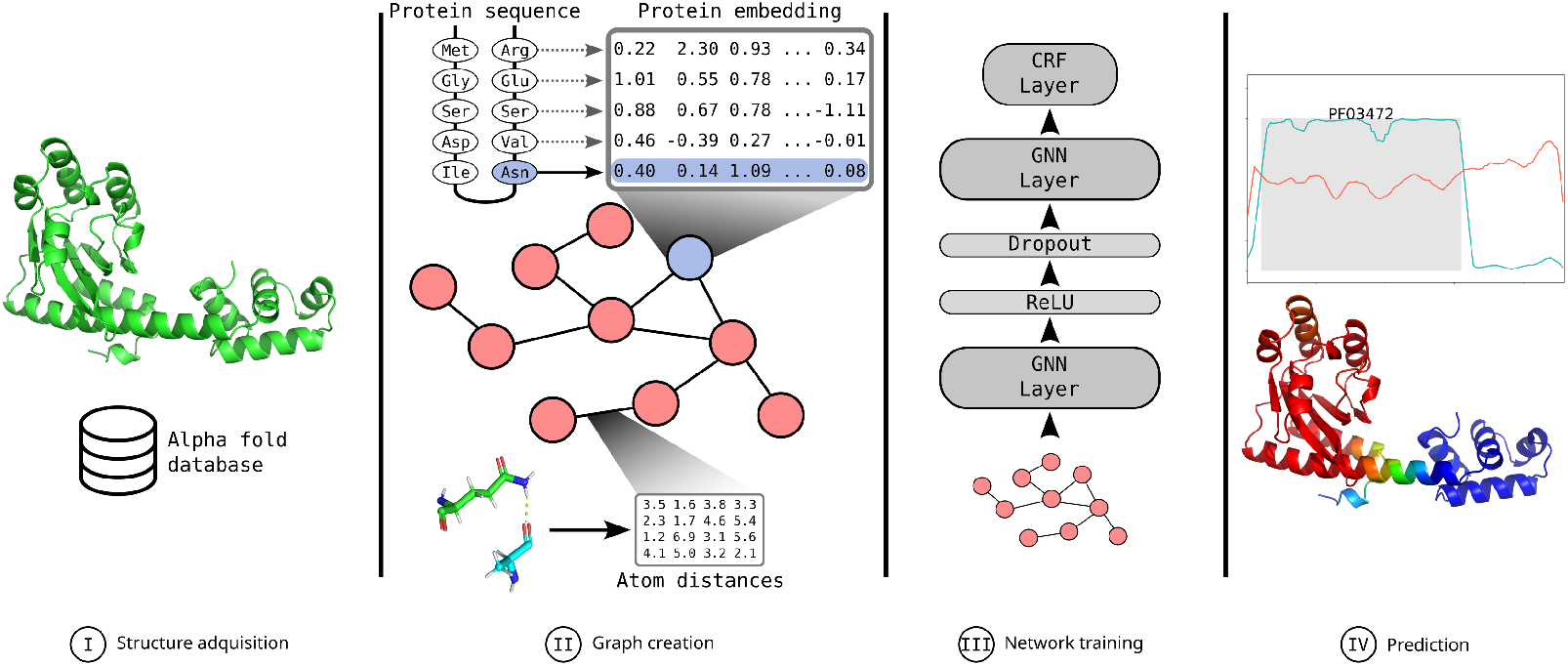
GNN2Pfam: novel pipeline for Pfam family annotation with a GNN model.

### Step I: Structure acquisition

First, for each raw protein sequence its corresponding 3D structure is obtained from the AlphaFold2 database in PDB format. Proteins absent from the database are dismissed. This procedure is performed in order to assure that every structure is complete and that all structures are consistent between them, thus avoiding possible errors or omissions from crystallographic measurements.

### Step II: Graph creation

For each amino acid in the input protein sequence, an embedding is obtained from a pLM; and at the same time its corresponding 3D structure of the protein (obtained in Step I) goes through the graph creation step. A graph is a data structure that models a set of relations (edges) between a set of entities (nodes); formally, *G* = (*V, E*), where *V* is the set of nodes and *E* is the set of edges. A bipartite graph is a graph composed of two disjoint sets of nodes, *V*_1_ and *V*_2_, such that every edge in the graph connects a node in *V*_1_ with a node in *V*_2_, formally *B* = (*V*_1_, *V*_2_; *E*). A protein 3D structure can be naturally represented as a graph *G*, where each amino acid residue is a node, thus the full sequence is *V*. Edges that connect nodes are the edge features *E*, here defined according to the spatial proximity of amino acids in the 3D structure: two residues are connected if the distance between them falls below a specified threshold. In our model, each node is assigned a vector corresponding to a pLLM embedding of the amino acid it represents, also capturing contextual information from the primary sequence.

Edge features are calculated for each connection between nodes and consist of a 4 × 4 matrix of the pairwise distances between the carbon, oxygen, nitrogen, and sulfur (CONS) atoms of the connected residues (see Figure 1, step II, below). Three different strategies were explored to define connections in the adjacency based on inter-residue distance: i) 10ÅC*β*: when the distance between beta carbon (C*β*) atoms is less than 10 Å; ii) 4ÅCe: when distance between residue centroids is less than 4 Å; and iii) 5ÅCONS: when the minimum distance between any pair of CONS atoms is less than 5 Å. The choice of threshold and reference atoms significantly affects graph density and neighborhood composition, which in turn impacts the information flow in the GNN layers. These different strategies resulted in graphs with varying connectivity patterns, as illustrated in Figure 2. Panel A) shows the protein A0A0A6P4U3_9GAMM and its corresponding AlphaFold2 structure ^1^. Figure 2, panel B) shows the connectivity map of the protein using the 10ÅC*β* criteria. The large radius used for this metric allows further amino acids to be considered as interacting pairs, which produces a densely connected graph. Figure 2, panel C) shows the connectivity using the more restrictive 4ÅCe criteria, where this shorter cutoff, combined with measuring from the amino acid’s center of mass, limits interactions to only very close residues, producing a sparse network. Figure 2, panel D) presents the 5ÅCONS metric, this being a middle point between the previous two criteria. It captures local residue environments effectively while minimizing the inclusion of spurious interactions. Among the tested approaches, the 5ÅCONS yielded the best performance in preliminary experiments with a small subset for domain prediction. This strategy likely provides a more detailed and functionally relevant view of residue interactions by capturing close-range physical contacts, regardless of backbone position or side chain type. Consequently, it was selected as the default adjacency criterion for the final GNN model.

**Figure 2:**
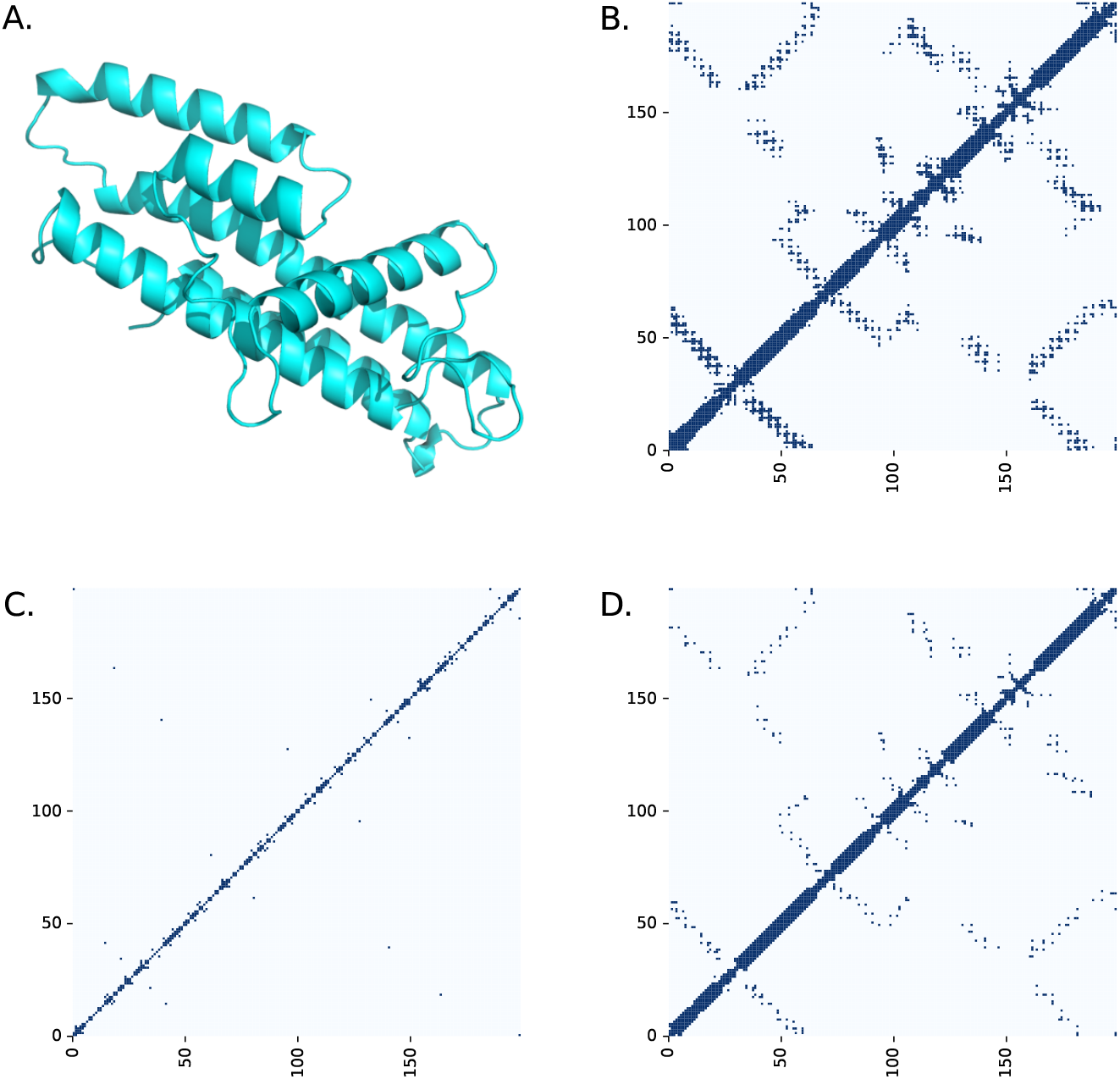
Examples of connectivity maps for the protein A0A0A6P4U3_9GAMM. A) AlphaFold2 3D structure. Connectivity maps: B) 10ÅC*β*, C) 4ÅCe, and D) 5ÅCONS.

### Step III: GNN2Pfam model training

At the next step of the pipeline, the GNN2Pfam model learns features hierarchically by iteratively aggregating information from the local neighborhood of each node. The GNN architecture has 2 Graph Attention Network (GAT) layers (25), which operate on graph-structured data, using masked self-attentional layers to improve the shortcomings of convolutions layers that can only represent grid-like structures. There is also a ReLU and a dropout layer. A GAT layer has the capability of specifying different weights to different nodes in a neighborhood, this way each node is able to attend to their neighborhood features without requiring costly matrix operations. To enhance sequence modeling and enforce structured predictions, the final layer of the model is a CRF layer (5), which is generally used for structured prediction. CRF can take context into account and predict a label for a single sample considering neighbouring samples as well. The predictions are then modelled as a graphical model, which represents the presence of dependencies between the predictions. This CRF layer captures dependencies between output labels, improving the consistency of predictions by considering contextual relationships rather than making independent predictions for each node. The model is trained using the Adam optimizer (14). A dropout layer is used during training to mitigate overfitting. The training objective is to minimize the cross-entropy loss between predicted and true Pfam domain labels.

### Step IV: Pfam family prediction at the output

At the output of the model, for each amino acid in the input protein sequence, a score is predicted for each Pfam family. It is important to highlight that at the output there is also an extra class named “no domain”, which models the cases where there is no reported Pfam family. That is, the output classes are all the possible Pfam families and the no domain class. The final predicted Pfam family can be calculated in three different ways: by maximum score, by coverage score, or by the area score. The maximum score is obtained by simply considering the highest score at the output along the protein, that is, the output class with the highest value along all the amino acids that belong to the test domain will be the predicted class. The coverage score considers how many times each class was the winner along all the amino acids of the protein sequence. Then the class with the largest coverage is considered the predicted class. Finally, the area score considers that all the scores of a class along the full sequence length constitute a curve and the area under that curve of scores is considered as the prediction score for the class. The output class with the largest area under the curve of scores is considered as the predicted one.

## 3. Materials and experimental setup

The sequences and Pfam annotations dataset used for training and testing this proposal is a subset from the data used in (2). Seed sequences from Pfam v.32.0 were split by clustering based on sequence identity. In order to assure remote homology (low similarity between training and testing sequences), single-linkage clustering at 25% identity within each family was used to build a clustered split. This provides a more realistic testing scenario, close to a real-world annotation problem, where testing sequences are not similar to the training ones. In our experiments we used the full proteins, not just the Pfam seed domain. For full protein representation, ESM2 was used, which provides embeddings of size 1,280 (20). The original dataset presented a high level of imbalance between partitions having, for example, families with all members in the test set, that is, with no representative members in the training set. Thus, we decided to use as a benchmark a subset of the data partitioning with a balanced number of Pfam families in train and test sets. This constitutes a sub-set of 6,169 proteins for training and 1,122 proteins for testing, belonging to 58 possible Pfam families, with a balanced number of examples both in train and test sets, keeping the same homology restrictions as the original dataset. The 3D structures of these proteins were obtained from the Alpha Fold database^2^, obtaining a final dataset of 7,291 proteins.

For the GNN model architecture, the following hyperparameters were explored (final model architecture values indicated in bold): representation layer size = 256, 512, **1**,**204**, 2,048, edge embedding size = **16**, number of layers = **2**, 4, 8, dropout = **0.1**, and learning rates = 1e-04, **1e-05**, 1e-06. We compared our proposed GNN2Pfam model with the HMM strategy currently used at Pfam for family annotation. To ensure a fair comparison, we trained the HMMs from scratch. We created custom MSAs using the sequences from the training set only, aligned with Muscle 3.8.31 (9). Using the MSAs generated we then employed HMMER 3.4 (8) to build the HMMs for each family domain and to make the predictions over the test set sequences. In order to analyse the model performance in a more challenging dataset, the benchmark was further divided to focus on families where at least one method had errors, either because it failed completely or because it confused the correct family with another one. The recall of the methods is calculated as 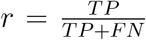, where TP is true positives, and FN is false negatives. The prediction error rate is calculated based on the recall as *e* = 1−*r*, by using the maximum score, the coverage score or the area score criteria.

## 4. Results

First, we performed an ablation study on the GNN2Pfam prediction model components. Table 1 shows the results on the ablation of the GNN and CRF component of the GNN2Pfam model proposed. The first column shows the mean errors (maximum, coverage and area) of the model when the CRF component is ablated. The second column shows the performance of the GNN when it has one GNN and one CRF layer. The last column shows the performance of the GNN when it has two GNNs and one CRF layer. The best (lowest) error results are obtained, in all cases, with the full model of 2 GNN+CRF layers. When the CRF component is ablated, performance is clearly affected, the errors achieved at the test partition are almost doubled.

**Table 1:** Ablation study on the elements of the GNN2Pfam model.

| **error** | **2 GNN** | **1 GNN+CRF** | **2 GNN+CRF** |
| --- | --- | --- | --- |
| **Maximum score** | 0.245 | 0.073 | **0.056** |
| **Coverage score** | 0.222 | 0.072 | **0.061** |
| **Area score** | 0.218 | 0.075 | **0.063** |

### 4.1 Pfam family prediction performance

Figure 3, Panel A) shows a violin plot with the median recall distribution on the full test partition of the proposed GNN2Pfam model versus the state-of-the-art HMM models used for Pfam families prediction, where the GNN2Pfam model clearly achieves the highest recall. For the GNN2Pfam and the HMM models in this test set, the median average recall is 1.0. However the median area error is 0.944 for GNN2Pfam while it is 0.815 for the HMMs. Figure 3, Panel B) depicts the detail of the recall for the families of the test partition where at least one of the models committed an error. That is, the set of Pfam families that were the hardest one to predict, for both, the state-of-the-art HMM models traditionally used by the Pfam database and also for GNN2Pfam. It can be seen here again that the GNN2Pfam model achieves the highest recall (1.0), that is, in spite of committing an error the GNN2Pfam recall is better than the HMMs models (0.75). Moreover, there is a statistically significant difference between these results according to a Mann-Whitney U test.

**Figure 3:**
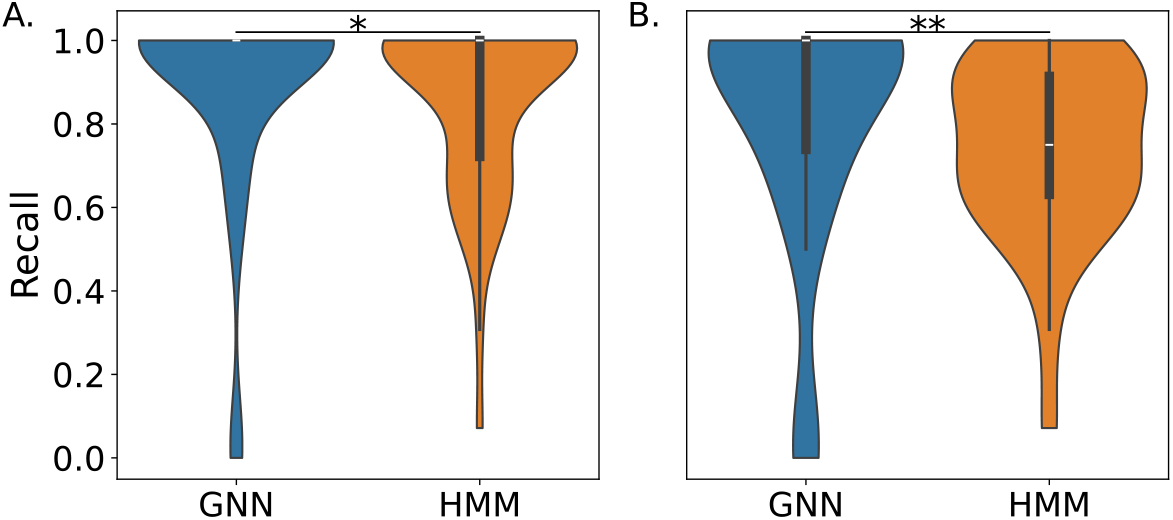
Violin plots of median recall distribution on the test partition of the proposed GNN2Pfam model versus the state-of-the-art HMM models used for Pfam families prediction. Panel A: full test partition. Panel B: Comparison of performance on the hardest testing cases: families of the test partition where at least one of the models commits an error. (*) 0.015 (**) 0.007 Mann-Whitney U test.

Figure 4 shows the details of the individual recall for the families of the test partition where at least one of the models has committed an error. This figure presents a more detailed analysis of results in Panel B in Figure 3. It can be seen that the GNN2Pfam model has the maximum recall, *r* = 1.0, in 56% of the cases, while providing a *r >* 0.50 in 90% of these hard test cases. The HMMs, instead, have lower recall and commit errors in most of the test cases. It is remarkable that in this hardest test the proposed GNN2Pfam model demonstrates better robustness. Notably, in the cases of the families that have very low recall according to the HMMs (PF03061 and PF01590), those are correctly predicted by the proposed GNN2Pfam model and with high recall.

**Figure 4:**
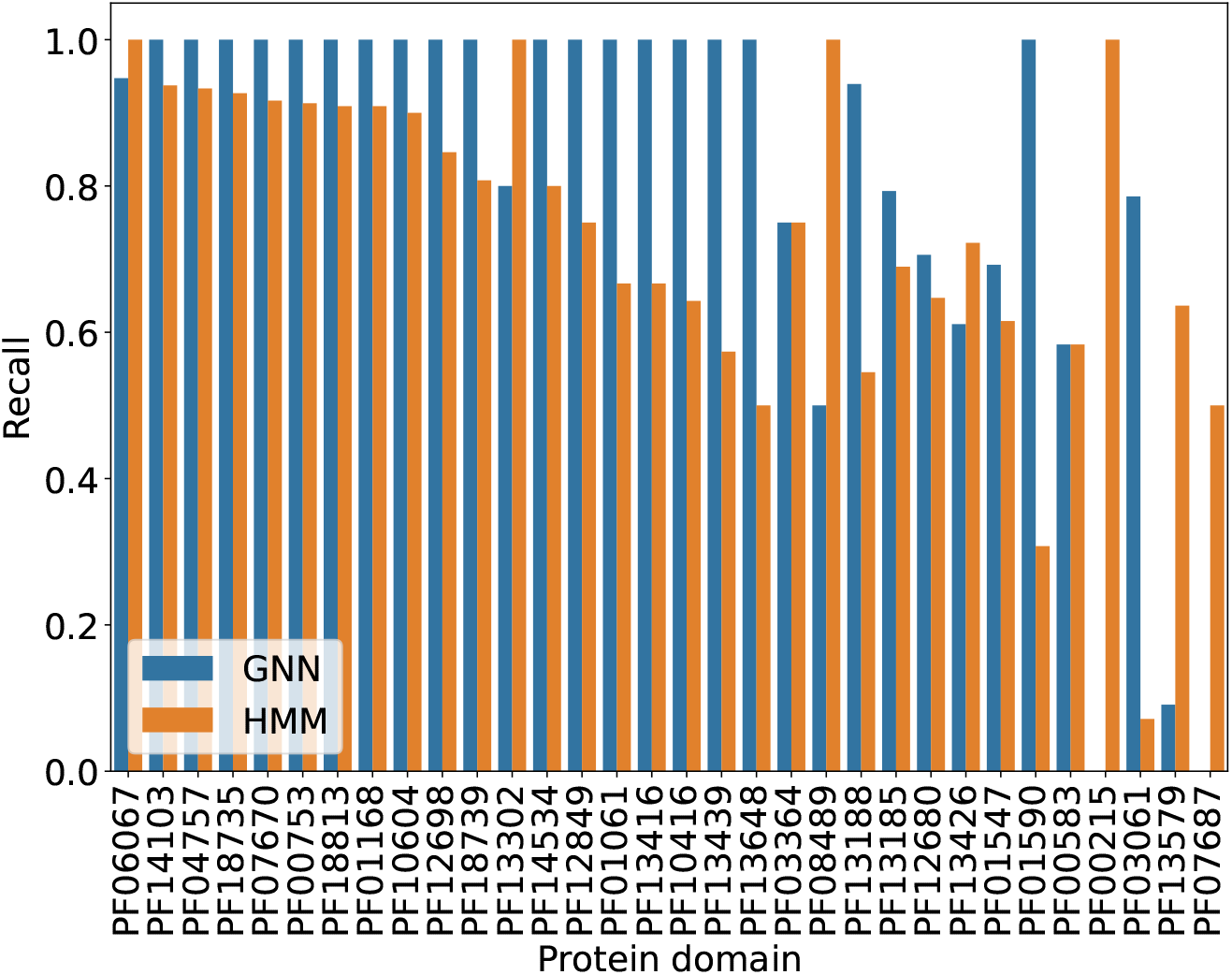
Detail on the per-family recall on the test partition of the proposed GNN2Pfam model versus the state-of-the-art HMM models used for Pfam families prediction, only for the families of the test partition where at least one of the models committed an error.

### 4.2 Pfam family prediction: some cases of study

In several cases, the GNN2Pfam model predicted the presence of a Pfam domain in a region of a protein where no domain annotation was present in the version of the Pfam database used for training (v32.0). Initially, these predictions were flagged as false positives. However, upon consulting the most recent release of Pfam, we found that in a number of these cases the domain predicted by the GNN2Pfam model is now annotated, suggesting that the model was able to anticipate correct annotations ahead of time.

For example, Figure 5 shows the GNN2Pfam output for the B9JSF3-_AGRVS protein. The x-axis indicates amino acid coordinates and the y-axis shows prediction scores. The gray shadow box marks the domain reported for this protein at Pfamv32.0 (PF03472), the green line shows the GNN2Pfam prediction for the output class PF03472, and the red line shows the “No domain” prediction along this protein. At the bottom, a heatmap presents the pLDDT (per-residue score of local confidence) value of each residue as reported by AlphaFold2, ranging from low confidence (yellow) to high confidence (blue). It can be seen that in this case the GNN2Pfam prediction has a perfect match with the beginning and ending parts labeled of the domain at the reference. This happens precisely at the intersection points of the GNN2Pfam model prediction curve (green) and the No domain curve (red).

**Figure 5:**
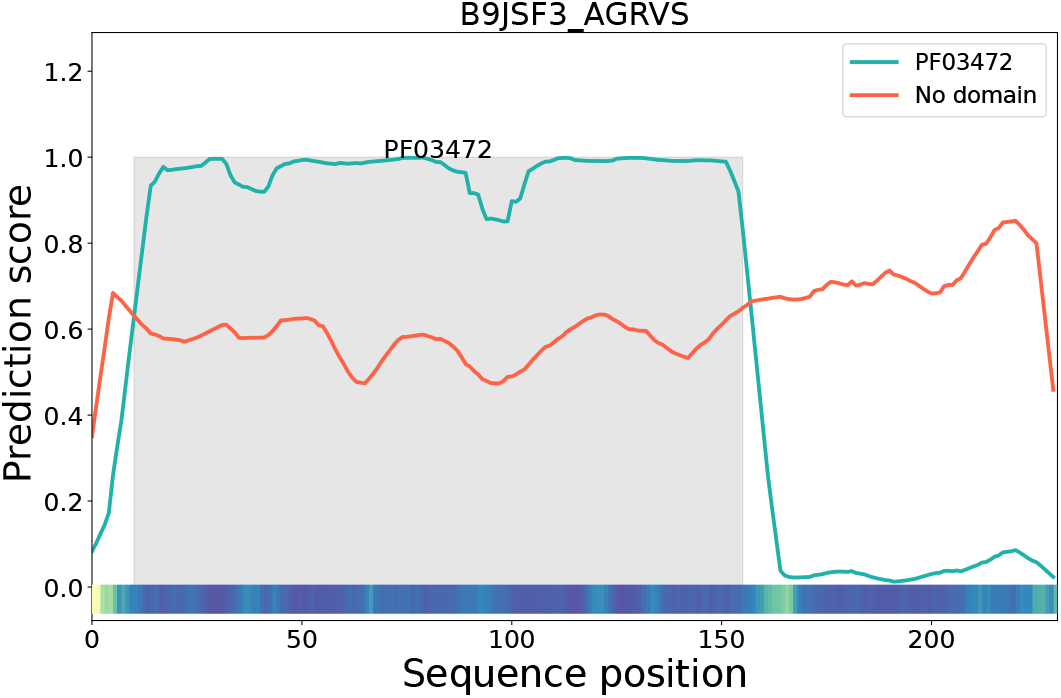
Example of GNN2Pfam output for the B9JSF3_AGRVS protein: GNN2Pfam prediction (green), No domain prediction (red), labeled domain at Pfamv32 (gray shadow box). Bottom: pLDDT value of each residue as reported by AlphaFold2.

The pLDDT values corresponding to this protein show that the corresponding 3D structure in the domain region has a high confidence, and also show low confidences precisely at the domain borders. All this information has been captured by the GNN2Pfam features, which have certainly helped the model to make a correct prediction.

Figure 6 shows another very interesting behavior of the GNN2Pfam model, where a “prediction error” according to the annotations in Pfam v32.0 was afterwards rectified in later versions of the database, in coincide with the GNN2Pfam model prediction. The figure shows in shaded gray, the Pfam v32.0 annotation for this protein (PF03364) from positions 9 to 138. However, the recent version of the annotation at Pfam v35.0 corrected it to be the domain PF10604 (light green box) between residues 4 and 149 for this protein. And it can be clearly seen that the GNN2Pfam model has actually predicted the PF10604 (green line) for this protein. This means that the GNN2Pfam model here had the capability of recognizing the pattern of the correct domain PF10604, in spite that this protein belonging to the test set was incorrectly annotated to another domain in the training set. It is also very interesting to note the AlphaFold2 annotation for PF10604 domain in this protein: where there is low confidence (yellow zone between positions 115 and 125), the GNN model had a drop in its scores that perfectly matches the low AlphaFold2 structural prediction confidence.

**Figure 6:**
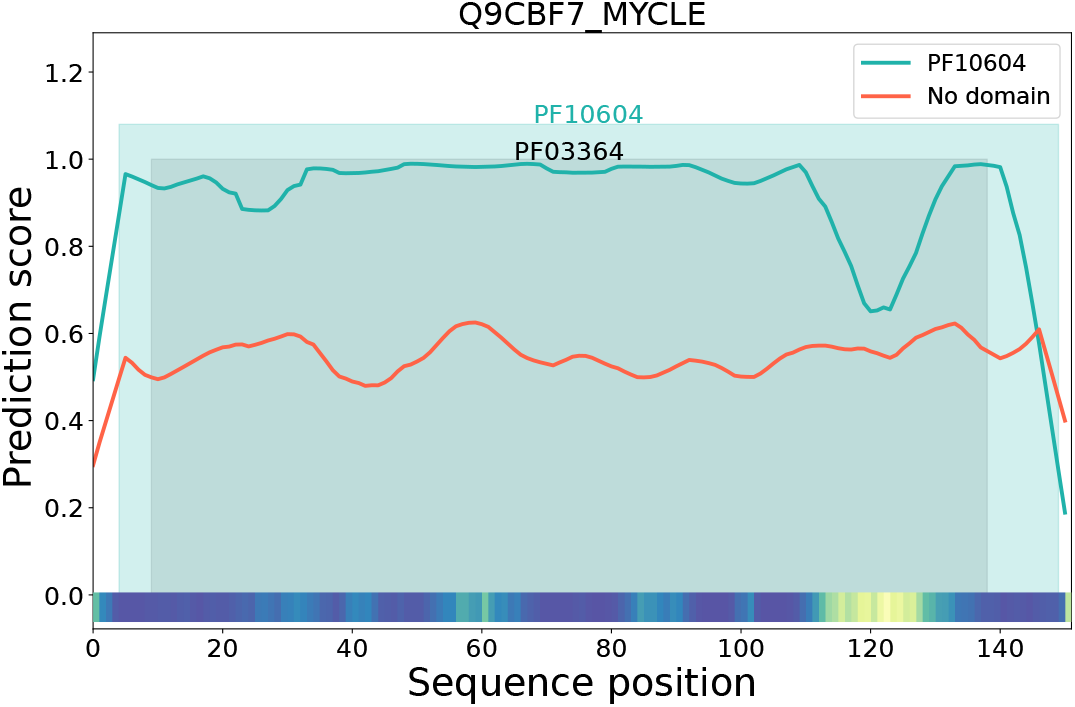
Example of a GNN2Pfam prediction for the protein Q9CBF7_MYCLE with the domain PF03364 labeled at Pfamv32.0 (gray box), but then corrected at the posterior version of Pfamv35.0 as domain PF10604 (light green box).

Figure 7 shows an example of a GNN prediction for the protein Q8YT62-_NOSS1 that has labeled the family PF13579 (gray box) according to Pfam v32.0. The proposed GNN2Pfam model predicts both the family PF13579: *Glycosyl transferase 4-like domain* (green line) and also the family PF13439: *Glycosyltransferase Family 4* (blue dotted line) with high scores. A deeper analysis reveals that both families are actually the same one. In fact, those have been integrated now at InterPRO with the code IPR028098 and both have the same description: *MshA belongs to the GT-B structural family of glycosyltransferases whose members have a two-domain structure with both domains exhibiting a Rossman-type fold. This entry represents the N-terminal domain found in MshA and the subfamily 4 of glycosyltransferases family 1*.” Interestingly, the No domain signal (red line) crosses at the green and dotted lines indicating beginning and ending domain coordinates.

**Figure 7:**
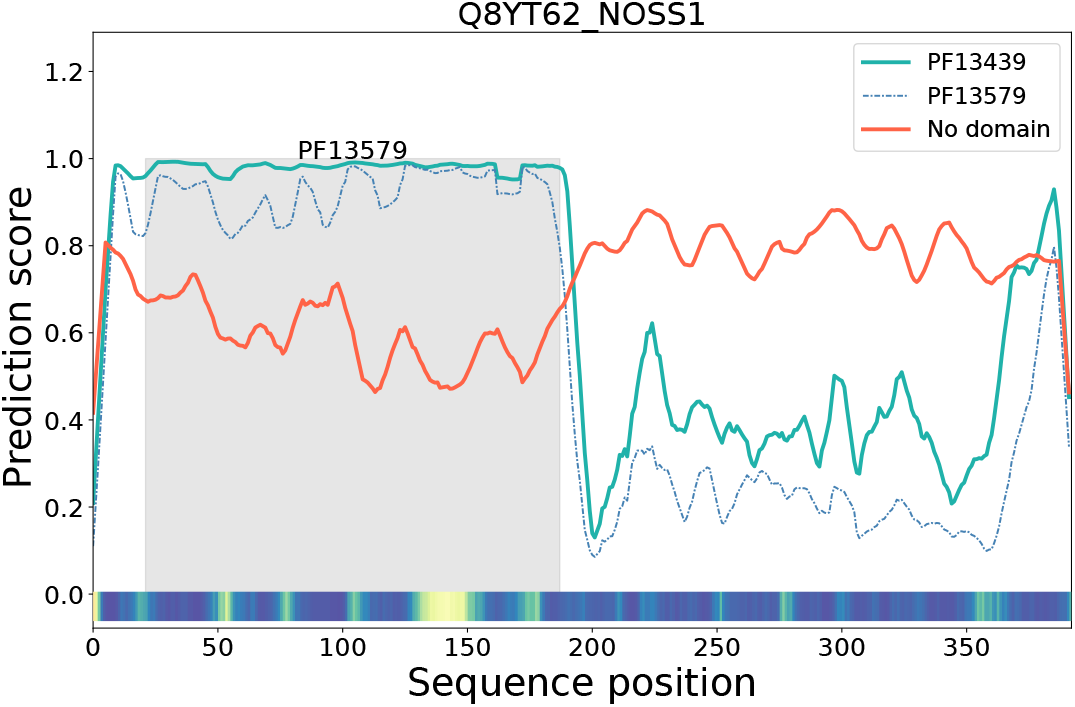
Example of a GNN2Pfam prediction for the protein Q8YT62_NOSS1 that, while Pfam v32.0 reports the family PF13579 as the correct label for the domain location (gray box), the GNN model predicts both the family PF13579 (green line) and also the family PF13449 (blue dotted line).

Figure 8 illustrates a case of structural instability detection by the GNN2-Pfam model. In several proteins, AlphaFold2-predicted structures are of low quality, which can negatively affect the models. This figure shows that regions where the prediction scores fall are aligned with the loss of AlphaFold2 confidence in the structure. That is, when there is low pLDDT AlphaFold2 confidence (yellow and orange regions, at the bottom), the GNN2Pfam output scores are low (around positions 50, 12, 170 and 260 in the sequence). At the correct domain location (gray box) with high AlphaFold2 score (blue), the GNN2Pfam scores for the PF01062 domain are high (green line) and intersect with the No domain curve at the beginning and ending points of the labeled domain. It is important to highlight that this result could not be achieved by simply having a threshold above a certain value (e.g. 0.5) for the No domain class. It is also very interesting to see here that the No domain signal, at the coordinates of no domain annotations (350 and up to the end of the sequence), shows a higher score than all the other classes that are incorrect, clearly indicating the No domain region.

**Figure 8:**
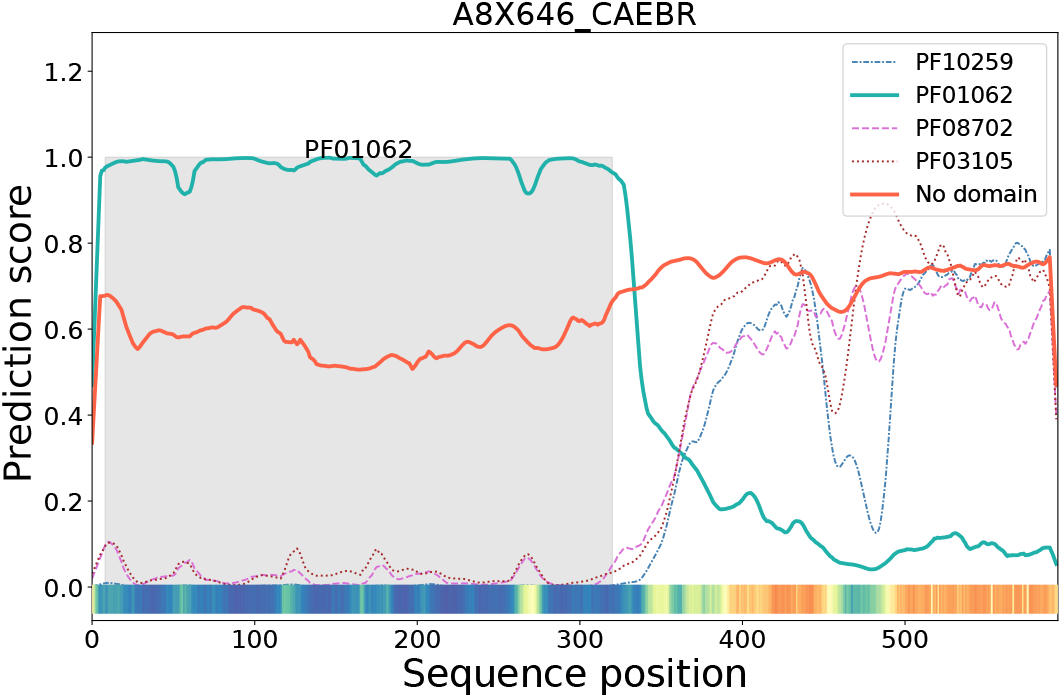
Example of low quality AlphaFold2-predicted structures that are detected by the GNN2Pfam model.

Finally, Figure 9 illustrates the most interesting application of the model for Pfam annotations: knowledge discovery of new locations of a domain. In Pfam v32.0 the protein D1B7T5_THEAS has only the PF13185 seed (GAF_2 domain) labeled between positions 175 and 313 (gray box). The output of the GNN2Pfam (green line) that crosses the No domain curve (red line) also shows two other locations of domains up and downstream of this protein sequence. Exactly in the same coordinates, GNN2Pfam also predicts the domain PF01590 (GAF domain). In the same version of Pfam v32.0 the downstream domain was annotated with the PF13185 domain by the HMMs as the GNN2Pfam model has effectively predicted. The domain located to the N terminal of the sequence remains yet unannotated by Pfam, but it is effectively identified by the InterPro database nowadays as IPR003018, which integrates both Pfam domains annotations PF13185 and PF01590. Thus, the knowledge discovered by the GNN2Pfam model has been effectively confirmed in a posterior version of the databases.

**Figure 9:**
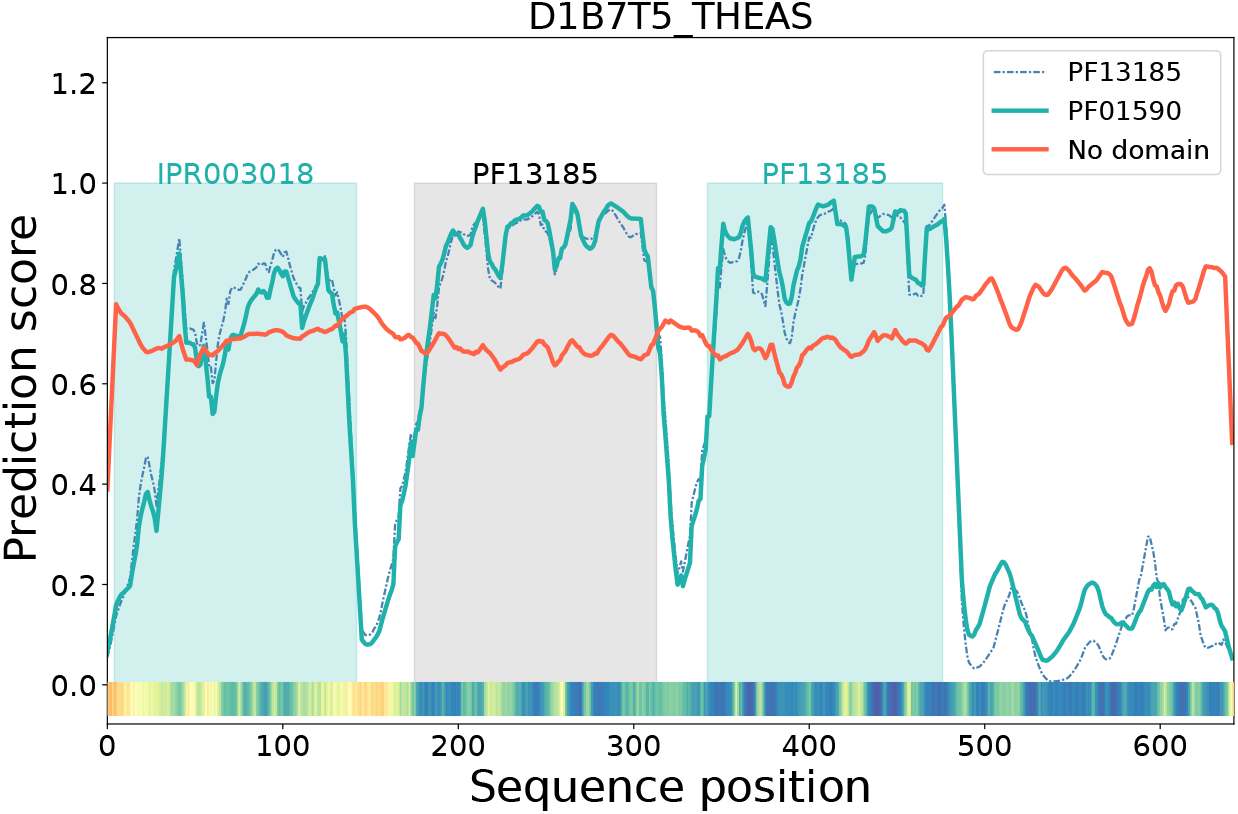
Example of knowledge discovery with the GNN2Pfam model and the protein D1B7T5_THEAS.

## 5. Conclusions

In this work we have proposed GNN2Pfam, a novel approach for Pfam family domain annotation with a graph neural network that integrates both sequence and 3D structure information into a novel architecture. Our proposal GNN2Pfam is the first domain prediction model integrating protein 3D structure together with sequence representation obtained from a pre-trained large language model. The results obtained indicate that the GNN2Pfam approach has largely overcome the state-of-the-art HMM used nowadays at Pfam, showing significantly better recall and lower error. It is very interesting to highlight also the potential of this model for knowledge discovered in yet annotated proteins, since in many cases the GNN2Pfam model predictions have been effectively confirmed in posterior versions of the Pfam database. These results suggest that GNN models, using both sequence embeddings from protein language models and 3D structure predictions, can be a key component for boosting future protein annotation tools. Future work involves extending the experimental setup with a larger database and newer versions of Pfam, as well as the automatic segmentation of domains, providing marks for beginning and end.

## Funding

This work was supported by ANPCyT PICT 2022 0086 and CAID-UNL 2024 0100097.

## Footnotes

1 https://www.uniprot.org/uniprotkb/A0A0A6P4U3/entry

2 https://alphafold.ebi.ac.uk/

## Notes

### Competing Interest Statement

The authors have declared no competing interest.

